# The developmental emergence of tonic and phasic REM sleep in rats

**DOI:** 10.1101/2025.11.26.690540

**Authors:** John P. Kobrossi, James C. Dooley

**Author notes:** Corresponding author: James C. Dooley, Ph.D. Lead Contact: James C. Dooley.

## Abstract

REM sleep is composed of two substates—phasic and tonic—that differ in their behavioral, sensory, and electrophysiological features. Although these substates are well characterized in adults, their developmental trajectory remains unclear. Here, we examined the development of tonic and phasic REM in rats from postnatal day (P) 12–24, spanning a period of rapid corticothalamic development. We recorded local field potentials and single units from primary motor cortex (M1), together with high-speed video and electromyographic recordings of the nuchal muscle. Periods of behavioral quiescence along with high delta power indicated NREM sleep, whereas periods of sustained muscle atonia and low delta power indicated REM sleep. At P16, M1 theta oscillations first appeared, and the delay to the first twitch increased, revealing the start of a distinct twitch-free portion of REM sleep. Motivated by this, we divided REM sleep into phasic and tonic periods, with and without twitching, respectively. Spiking activity and gamma power were consistently higher during phasic REM. At P20, phasic REM also showed faster theta oscillations than tonic REM. At P24, tonic REM was accompanied by a distinct alpha oscillation. These results show that the features distinguishing the two REM substates appear sequentially across development, revealing a progressive differentiation of REM sleep into tonic and phasic periods, a developmental refinement that may support increasingly complex forms of sleep-dependent plasticity.

**Significance Statement:** Infancy is marked by rapid circuit formation and by the dominance of REM sleep, a state thought to support early neural development. Yet sleep itself undergoes massive changes throughout this period, and the developmental timeline of REM sleep remains poorly understood. Using neural recordings and high-speed video from developing infant rats, we show that REM sleep gradually divides into two substates—tonic and phasic—across infancy. These substates exhibit age-dependent differences in movement, neural firing, and cortical oscillations, revealing increasing microstructural complexity in REM sleep. Our findings identify when these substates first emerge and how their defining features unfold over time, providing a framework for understanding how REM sleep supports developmental plasticity throughout early life.

## Introduction

Since its discovery in humans, rapid eye movement (REM) sleep has been described as a paradoxical state, combining wake-like brain activity with sleep-like behavior. This paradox becomes more striking when REM sleep is examined at finer resolution. In adults, REM sleep alternates between phasic and tonic periods, each marked by distinct patterns of movement, cortical activity, and sensory responsiveness (Ermis et al., 2010; Simor et al., 2017, 2020). In humans, phasic REM features bursts of eye movements and muscle twitches against a background of atonia and is accompanied by elevated gamma power and high arousal thresholds (Datta and O’Malley, 2013). Tonic REM, by contrast, is defined by behavioral quiescence, stronger alpha/beta activity, and lower arousal thresholds (Waterman et al., 1993; Wehrle et al., 2007). Despite growing appreciation of this microarchitecture, remarkably little is known about when in development these distinct REM substates first emerge.

In early postnatal life, research in humans and rodents alike shows that the majority of time is spent in REM sleep, also called active sleep in infancy (Jouvet-Mounier et al., 1969; Knoop et al., 2021). Throughout infancy, the twitches of REM sleep drive sensory feedback in the developing nervous system (Khazipov et al., 2004; Sokoloff et al., 2015; Dooley and Blumberg, 2018; Dooley et al., 2020). These twitches are abundant at birth, but decline sharply over the first few postnatal weeks (Blumberg et al., 2005; Reid et al., 2025). During this same developmental window, thalamocortical networks undergo rapid development: delta rhythms are established during NREM sleep (Seelke and Blumberg, 2008), sensory responses become more temporally precise (Dooley and Blumberg, 2018; Dooley et al., 2021), and theta oscillations emerge in the brainstem and hippocampus (Del Rio-Bermudez et al., 2017; Muessig et al., 2019). Thus, late infancy (P12–P24) represents a window in which REM sleep physiology and cortical function are in continuous flux.

Tonic and phasic REM have been described in adult rodents (Brankačk et al., 2012; Meng et al., 2021; Dong et al., 2022; Bueno-Junior et al., 2023), with each substate sharing many of the features described in humans (see **Table 1**). Although prior research has described age-related changes in the proportion of tonic and phasic REM sleep across development (Vogel et al., 2000), this work did not explore whether there are differences in cortical oscillations or neural activity across age. Consequently, it remains unclear whether these substates emerge fully formed or whether their constituent features develop gradually. Characterizing this developmental progression is essential for linking the well-characterized behavioral and electrophysiological features of REM sleep in infancy to the microarchitecture of REM sleep in adults.

**Table 1.** Summary of reported differences between tonic and phasic REM sleep in human and mouse studies. This table compiles published comparisons of tonic versus phasic REM across species, highlighting differences in arousal threshold and oscillatory activity. Arousal threshold differences were assessed using auditory stimuli and are indicated by the method used: ERP (event-related potentials), BOLD (fMRI blood-oxygen-level–dependent responses), or Behav. (behavioral measures such as awakening or suppression of eye movements). For each frequency band, an X denotes a significant difference between substates, a + denotes a trend or partial effect, and blank cells indicate no reported difference. Asterisks mark cases where the analyzed frequency band deviated notably from the canonical ranges listed in the header (except for gamma, which commonly represents a narrower sub-range within the broader 30–200 Hz band). A dash (-) indicates that arousal threshold was not tested. ^✝^Substate sensory threshold data could be inferred indirectly from patch-clamp data but were not explicitly quantified. ^‡^These authors described the two REM substates as “REM with whisking” and “REM without whisking,” which correspond to phasic- and tonic-like periods, respectively.

| Citation | Species | Dev. Age | Sensory Thresh. | Delta 1-4 Hz | Theta 4-8 Hz | Alpha/Beta 8-30 Hz | Gamma 30-200 Hz | Notes |
| --- | --- | --- | --- | --- | --- | --- | --- | --- |
| (Sallinen et al., 1996) | Human | Adult | ERP |  |  |  |  |  |
| (Takahara et al., 2006) | Human | Adult | ERP |  |  |  |  |  |
| (Wehrle et al., 2007) | Human | Adult | BOLD |  |  |  |  |  |
| (Price & Kremen, 1980) | Human | Adult | Behav. |  |  |  |  |  |
| (Takahara et al., 2002) | Human | Adult | ERP |  |  |  |  |  |
| (Koroma et al., 2020) | Human | Adult | ERP |  |  |  |  |  |
| (Stuart & Conduit, 2009) | Human | Adult | Behav. |  |  |  |  |  |
| (Ermis et al., 2010) | Human | Adult | Behav. | X | X | X | X |  |
| (Avigdor et al., 2025) | Human | Adult | - | X | X |  | X |  |
| (De Carli et al., 2016) | Human | Adult | - |  |  | X |  |  |
| (Waterman et al., 1993) | Human | Adult | - |  |  | X |  |  |
| (Frauscher et al., 2016) | Human | Adult | - | X | X | X |  |  |
| (Jouny et al., 2000) | Human | Adult | - | X | X | X | X |  |
| (Nishida et al., 2005) | Human | Adult | - |  |  | X | + |  |
| (Gross & Gotman, 1999) | Human | Adult | - |  |  | + | X |  |
| (Simor et al., 2019) | Human | Adult | - | X* |  | X | X | *3-4 Hz |
| (Whitehead et al., 2019) | Human | Neonates | - | X | X |  |  |  |
| (Simor et al., 2016) | Human | 4-8 years | - | X | X | X | X |  |
| (Boscher et al., 2024)* | Mouse | Adult | † | X* |  | X |  | *2-6 Hz |
| (Brankačk et al., 2012) | Mouse | Adult | - |  | X |  | X |  |
| (Meng et al., 2021) | Mouse | Adult | - |  | X | X |  |  |
| (Dong et al., 2022)* | Mouse | Adult | - |  | X |  |  |  |
| (Bueno-Junior et al., 2023) | Mouse | Adult | - |  | X | X |  |  |

Here, we recorded local field potentials and single-unit activity from the forelimb region of primary motor cortex (M1) in unanesthetized, head-fixed P12–P24 rats. Behavioral states were characterized via continuous monitoring of nuchal EMG and high-speed video. REM-sleep-associated theta oscillations first appeared in M1 at P16. At the same time, the interval from REM onset to the first twitch lengthened, revealing sustained periods of REM sleep without twitches. Phasic REM consistently showed stronger gamma power and increased neural activity, but across P16 and P24, additional components developed that further differentiated REM sleep into tonic and phasic substates. By P20, theta oscillations during phasic REM were significantly faster than tonic REM, and the duration of tonic REM bouts increased. At P24, tonic REM was also accompanied by a distinct fast-alpha oscillation. Notably, the rate of twitches remained stable within phasic bouts across ages, even as the overall number of twitches declined. Thus, REM sleep continues to develop across infancy—from an early, largely undifferentiated state to one composed of tonic and phasic substates, each with distinct oscillatory signatures.

## Materials and Methods

### Experimental model and subject details

For this experiment, we recorded neural activity from the primary motor cortex (M1) in Sprague–Dawley rats (*Rattus norvegicus*) in four age groups: postnatal day (P) 12 (N = 9 pups; 30.5 ± 1.9 g; 3 male), P15–16 (hereafter referred to as P16; N = 11 pups; 43.8 ± 3.1 g; 8 male), P18–20 (P20; N = 12 pups; 56.1 ± 4.4 g; 7 male), and P22–24 (P24; N = 10 pups; 68.9 ± 4.4 g; 6 male). Each age cohort consisted of animals from separate litters. Although this study centers on M1 activity, recordings were obtained from the forelimb representation of M1 in combination with either the ventrolateral or ventroposterior nucleus of the thalamus. A subset of these animals has been include in prior studies that characterize their movement-related activity (see Dooley et al., 2021; Reid et al., 2025).

Pups were reared with their dam in standard laboratory cages (48 × 20 × 26 cm) maintained under controlled temperature and humidity on a 12:12 h light–dark cycle, with unrestricted access to food and water. The day of birth was designated P0, and litters were reduced to eight pups by P3. Each pup had at least four littermates until P12 and a minimum of two littermates on the day of recording. Animals remained with the dam until testing, and no pups were weaned before data collection. All procedures followed the National Institutes of Health *Guide for the Care and Use of Laboratory Animals* (NIH Publication No. 80–23) and were approved by the Institutional Animal Care and Use Committees of the University of Iowa and Purdue University.

### Method Details

*Surgery.* A pup of appropriate body weight and, for P12 animals, with a visible milk band was selected from the litter and anesthetized using isoflurane (3–5%; Phoenix Pharmaceuticals, Burlingame, CA). After shaving the scalp, care was taken to preserve all vibrissae. To monitor behavioral state, pairs of custom bipolar hook electrodes (0.002-inch diameter, epoxy-coated wire; California Fine Wire, Grover Beach, CA) were implanted into the nuchal and contralateral biceps muscles. Carprofen (5 mg/kg, subcutaneous; Putney, Portland, ME) was administered for postoperative analgesia.

The scalp was then excised, and a topical anesthetic (bupivacaine; Pfizer, New York, NY) was applied to the exposed skull. After cleaning and drying the skull surface with a mild bleach solution, small areas of residual bleeding were cauterized as needed in P20 and P24 pups to ensure complete dryness of the skull prior to headplate attachment. The surrounding skin was sealed with Vetbond (3M, Minneapolis, MN), and a custom stainless-steel headplate (Neurotar, Helsinki, Finland) was affixed to the skull using cyanoacrylate adhesive.

A 1.8-mm trephine drill (Fine Science Tools, Foster City, CA) was used to open a craniotomy above the forelimb area of M1 (0.5 mm anterior, 2.2–2.5 mm lateral to bregma). When additional recordings were made, a second craniotomy was placed above the thalamus (2.0–2.8 mm caudal to bregma, 2.2–2.5 mm lateral) as described previously (Dooley et al., 2021).

Following surgery, pups were positioned in a custom head-fixation frame mounted to a Mobile HomeCage (NTR000289-01; Neurotar, Helsinki, Finland). The height of the head above the cage floor was adjusted according to age such that the body rested naturally on the elbows during REM-related atonia (P12: 35 mm; P16: 38 mm; P20–P24: 40 mm). EMG leads were secured along the back to prevent tangling. Pups recovered from anesthesia within 15 minutes and were allowed 1–2.5 hours to acclimate to the head-fixation apparatus. This period allowed both electrode stabilization and the reestablishment of normal neural activity (Domínguez et al., 2021). At the start of recording, all animals displayed characteristic sleep–wake behaviors, including twitching, grooming, and spontaneous locomotion.

### Recording Environment

Recordings took place inside a sound-attenuating Faraday cage illuminated with constant, flicker-free red light (630 nm; Waveform Lighting, Vancouver, WA). A continuous broadband noise (70 dB) was played throughout the session to mask environmental sounds. To promote extended periods of REM sleep, ambient temperature was maintained between 26.5 °C and 29 °C (Szymusiak and Satinoff, 1981). The pup’s head was oriented away from the room entrance to minimize visual disturbance. All monitoring and data collection were performed remotely, with the experimenter positioned outside the room to reduce environmental disruptions and maintain a calm recording environment for the animal.

### Electrophysiological Recordings

Nuchal and biceps EMG leads were connected to the analog inputs of a Lab Rat LR-10 system (Tucker-Davis Technologies, Gainesville, FL). EMG signals were amplified, digitized at ∼1.5 kHz, and high-pass filtered at 300 Hz.

Prior to implantation, 16-channel silicon depth electrodes (A1×16-3mm-100-177-A16) were coated with the fluorescent tracer DiI (Vybrant DiI Cell-Labeling Solution; Life Technologies, Grand Island, NY). Electrodes were lowered into the forelimb representation of M1 using a multiprobe micromanipulator (New Scale Technologies, Victor, NY) operated via an Xbox controller (Microsoft, Redmond, WA). The array was advanced until all sites were located beneath the dura, typically at a depth of ∼1500 µm. A chlorinated Ag/Ag-Cl wire (0.25 mm diameter; Medwire, Mt. Vernon, NY) inserted into the contralateral occipital cortex served as the common reference and ground. Neural activity was digitized at ∼25 kHz with a 0.1 Hz high-pass filter to reduce signal drift and notch filters at 60, 120, and 180 Hz.

Recordings were collected continuously for 3–6 hours using SynapseLite software (Tucker-Davis Technologies). The Mobile HomeCage system allowed stable head fixation while permitting natural behaviors—including locomotion, grooming, and sleep—throughout the session.

### Video Collection and Synchronization

Video recordings were temporally aligned with the electrophysiological data to permit quantification of movement and behavioral state, as described previously (Dooley et al., 2020). Each pup was enclosed within a transparent chamber positioned inside the Mobile HomeCage, allowing full visual access during recording. A single Blackfly-S camera (FLIR Integrated Systems, Wilsonville, OR) was mounted at a 45° angle relative to the animal’s head, centered on the right forelimb contralateral to the cortical recording site. This setup provided a clear view of the right whiskers, forelimb, hindlimb, and tail. Video was captured in SpinView (FLIR Integrated Systems) at 100 frames per second, with a 7-ms exposure time and a resolution of 720 × 540 pixels.

To synchronize video and electrophysiological recordings, a red LED positioned within the camera’s field of view was driven by SynapseLite (Tucker-Davis Technologies) to emit a 100-ms pulse every 3 seconds. The timing of these flashes served as a reference signal for frame alignment. Custom MATLAB scripts quantified the number of frames between successive LED pulses to verify recording stability and detect dropped frames. In the rare event that the expected interval (300 frames, corresponding to the 3-s period at 100 frames/s) was not met, a placeholder frame was inserted at the missing location. This procedure maintained frame-by-frame correspondence between the video and electrophysiological data, ensuring synchronization accuracy within one frame (10 ms) across the entire recording session.

### Histology

Following completion of the recording session, pups were deeply anesthetized with ketamine/xylazine (10:1; >0.08 mg/kg) and transcardially perfused with 0.1 M phosphate-buffered saline (PBS), followed by 4% paraformaldehyde (PFA). Brains were removed, post-fixed in 4% PFA for at least 24 hours, and then transferred to a 30% sucrose solution for a minimum of 24 hours prior to sectioning.

To verify electrode placement within the M1 forelimb representation, cortices were sectioned either tangentially to the pial surface or in the coronal plane. For tangential preparations, the right hemisphere was carefully separated from the underlying subcortical tissue and flattened between two glass slides spaced 1.5 mm apart for 5–15 minutes, with light pressure applied using 10 g weights. All brains were cut at a thickness of 80 µm. Wet-mounted sections were imaged under a fluorescent microscope (Leica Microsystems, Buffalo Grove, IL) at 2.5x magnification to visualize DiI fluorescence and confirm electrode tract locations.

Cortical tissue was stained for cytochrome oxidase (CO) to delineate primary cortical areas, which are well defined by this method during the developmental period studied (Seelke et al., 2012). The staining solution consisted of cytochrome C (3 mg/10 mL; Sigma-Aldrich), catalase (2 mg/10 mL; Sigma-Aldrich), and 3,3′-diaminobenzidine tetrahydrochloride (DAB; 5 mg/10 mL; Spectrum, Henderson, NV) dissolved in equal parts phosphate buffer and distilled water. Sections were incubated on a shaker at 35–40°C (approximately 100 rpm) until the tissue was well differentiated (approximately 3-6 hours), then rinsed in PBS, mounted on glass slides, and air-dried for at least 48 hours. Once dry, slides were cleared in citrus clearing solvent (Richard-Allan Scientific) and cover-slipped with DPX.

Final images were collected at 2.5x or 5x magnification. When needed, multiple images were stitched into a single composite using Microsoft Image Composite Editor (Microsoft, Redmond, WA), allowing electrode tracks to be visualized relative to areal and laminar boundaries of cortex.

### Local Field Potential

For analysis of local field potentials (LFP), raw neural signals were first smoothed with a Gaussian moving kernel (half-width: 0.5 ms) to reduce high-frequency noise and then resampled to approximately 1 kHz.

For each recording, one channel was selected for LFP analyses using the following procedure. Channels exhibiting excessive noise or poor correlation with neighboring sites were first excluded. For the remaining channels, a zero-phase digital bandpass filter was applied to isolate delta (0.5–4 Hz) frequency bands. A trained scorer then identified at least five NREM sleep epochs, each lasting a minimum of 10 seconds. The channel displaying the highest delta power during these NREM segments was chosen for all subsequent LFP analyses. This approach typically selected more superficial cortical sites (at a depth of 200-400 µm, as delta power generally declined with depth along the electrode shank. In contrast, theta activity during REM sleep was generally uniform across channels and cortical layers.

### ROI Movement Classification

To quantify movement events, frame-by-frame changes in pixel intensity were analyzed within predefined regions of interest (ROIs) using custom MATLAB scripts(Dooley et al., 2021). For each ROI, the number of pixels exhibiting an intensity change greater than 5% between consecutive frames was summed, yielding a continuous 100 Hz signal reflecting movement-related pixel variation. This automated detection procedure identified candidate movements—either twitches or wake movements—that were subsequently verified by visual inspection of synchronized video recordings to confirm both the occurrence of movement and the specific body part involved.

For twitch analysis, the ROI corresponding to the relevant body part (e.g., forelimb) was examined in Spike2 (Version 8; Cambridge Electronic Design, Cambridge, UK) alongside the video. Peaks in the ROI-derived time series were validated as discrete twitches. Twitch onset was defined as the first frame in which movement activity was detected. For the limbs and tail, this approach reliably captured every visible twitch, as even closely spaced events exhibited distinct onsets and offsets. In contrast, whisker movements sometimes consisted of rapid alternating protractions and retractions without clear boundaries, in which case only the first twitch in a series was counted.

Wake movements were assessed using a ROI encompassing the entire animal because body-part-specific ROIs were less reliable given the variability in limb position during wakefulness. Only wake movements involving the forelimb were included in subsequent analyses. The onset of each wake movement was defined as the first frame in which the forelimb moved independently. To ensure that labeled wake events represented distinct transitions from inactivity to movement, only the initial movement in a bout was marked, and additional movements were annotated only if separated by at least 500 ms of behavioral quiescence.

### Classification of Behavioral State

Behavioral state was classified based on concurrent analysis of cortical delta activity and nuchal EMG. For each pup, the rectified EMG signal was dichotomized into periods of high muscle tone and atonia. Five representative segments of each condition were selected (at least 10 seconds in duration), and their median amplitudes were computed. The midpoint between these two medians served as the threshold for distinguishing tone from atonia. Using a running 1-s average of the rectified EMG, values exceeding this threshold were classified as periods of tone, whereas values below the threshold were classified as atonia.

A similar procedure was applied to the LFP signal to differentiate high-delta and low-delta periods. Segments of NREM sleep and REM sleep (defined below) were used to establish median amplitudes, and the midpoint between them was taken as the delta threshold.

Periods of high delta power and behavioral quiescence were classified as NREM sleep. Periods of nuchal atonia and low delta power were classified as REM sleep. To avoid spurious state transitions, any putative change in state was required to persist for at least 5 s; otherwise, the animal was considered to have remained in the prior state. Data were analyzed continuously (i.e., not in 5 second bins), allowing bouts to begin at any time, but transient deviations shorter than 5 seconds were ignored. Thus, brief arousals (including postural adjustments and stretching) lasting less than 5 seconds in duration did not end a NREM bout. Periods not classified as NREM or REM sleep were considered wake. Active wake was defined as that part of the wake period that was within 3 s of movement detected by the ROI Movement Classification; all other timepoints were considered quiet wake. This ensured that all quiet wake periods were not during sleep, and at least 3 s from any movement.

### Division of tonic and phasic REM substates

Within REM sleep, periods were further classified as tonic or phasic REM based on the occurrence of myoclonic twitches. Twitch events were identified from the manually scored movements of all body parts (forelimb, hindlimb, whisker, and tail). For each twitch, the window centered on the event (±2 s from twitch onset) was designated as phasic REM. All remaining periods of REM sleep that did not overlap with any phasic window were classified as tonic REM. This approach ensured that phasic REM encompassed both the twitch itself and any associated oscillatory change preceding or following the movement, while tonic REM reflected periods of muscle atonia and relative motor quiescence.

### Power Spectrum Analysis

LFP activity was analyzed across behavioral states, including quiet wake, active wake, REM sleep, and NREM sleep. Power spectral densities were computed in MATLAB using the pspectrum() function with default parameters. The resulting spectra were expressed relative to quiet wake by converting power values to decibel (dB) units. Quiet wake served as the normalization baseline because it occurs alongside behavioral quiescence, removing the possibility of movement artifact, and lacks strong state-dependent rhythmic activity—such as the theta oscillations typical of REM sleep or the delta and sleep spindle activity characteristic of NREM sleep.

### Quantification and statistical Analysis

#### Spike Sorting

Neural recordings acquired in SynapseLite were converted to binary format using custom MATLAB scripts and processed in Kilosort 2.0 (Pachitariu et al., 2016). Prior to spike detection, data were whitened (covariance-standardized) and band-pass filtered between 300 and 5000 Hz. Event waveforms were sorted into clusters through template matching. The initial spike detection threshold was set to six standard deviations below the mean signal, followed by a second-pass threshold of five standard deviations. A minimum firing rate of 0.01 spikes/s was required for inclusion, and the bin size for template estimation was set to 262,400 samples (approximately 11 s). All other Kilosort parameters were maintained at default values.

Cluster visualization and manual curation were performed in Phy2 (Rossant and Harris, 2013). Units were classified as putative single neurons if their spike waveforms consistently conformed to a single template, appeared as distinct, well-isolated clusters in principal component space, and exhibited a clear refractory period in their auto-correlogram (manifested as reduced firing at zero-time lag). Clusters meeting the first two criteria but lacking a refractory period were categorized as multi-units and excluded from further analysis.

Units displaying waveform characteristics suggestive of electrical noise, firing rates below 0.01 spikes/s, or amplitude instability over time were also excluded. To verify unit isolation, cross-correlograms of all single- and multi-unit pairs recorded on neighboring channels were compared. When cross- and auto-correlograms exhibited similar features, clusters were merged and reclassified as multi-units when appropriate.

#### Statistical Analyses

All statistical analyses were conducted in MATLAB. The significance threshold (α) was set at 0.05 for all tests, and Bonferroni corrections were applied when multiple comparisons were performed. Unless otherwise stated, data are presented as mean ± standard error of the mean (SEM). Depending on the experimental design, differences were evaluated using a one-way ANOVA, two-way ANOVA, or *t*-tests.

Violin plots were generated with outliers removed, as determined by MATLAB’s isoutlier() function. This function identifies and excludes data points that deviate by more than three scaled median absolute deviations from the sample median. The survivor plot of tonic REM bout length was tested for significance using the MatSurv function for MATLAB with its default parameters (Creed et al., 2020).

## Results

To establish the developmental emergence of tonic and phasic REM sleep, we recorded single-unit activity in unanesthetized rats at P12, P16, P20, and P24. Rats were tested in the Mobile HomeCage, an air table that allows rats to locomote while head-fixed, in a recording environment that was approximately 29° C, which is permissive of REM sleep (Szymusiak and Satinoff, 1981). Together, this allowed us to record significant periods of REM and NREM sleep on the day of surgery. Neurophysiological signals were recorded in the forelimb region of M1 (P12: N = 9 pups; P16: N = 11 pups; P20: N = 12 pups; P24: N = 10 pups). Neural activity, electromyographic activity of the nuchal and biceps muscles, and high-speed video (100 frames/s) were recorded continuously for 3-6 h. Recordings were performed entirely during the lights-on period (1000 to 1800 hours).

## REM and NREM sleep oscillations from P12 to P24

In rodents, myoclonic twitches of the skeletal muscles often define phasic REM, whereas periods of REM sleep lacking twitches defined as tonic REM (Vogel et al., 2000; Brankačk et al., 2012; Meng et al., 2021; Dong et al., 2022; Bueno-Junior et al., 2023). Thus, to be inclusive of both tonic and phasic REM, we avoided defining REM based on twitches and instead relied on two electrophysiological criteria. The first criterion was nuchal muscle atonia. Because nuchal atonia may also occur during NREM sleep (Seelke and Blumberg, 2008), the second electrophysiological criteria was low delta power. The change in nuchal tone at the onset and offset of REM sleep, along with the decrease in delta power at the onset of REM sleep, are shown in **Figure 1**. Periods of behavioral quiescence and high delta power were classified as NREM sleep. All timepoints not classified as REM or NREM sleep were classified as wake, which we split into periods of quiet wake (lacking movement) or active wake (with movement). Rats cycled between NREM sleep, REM sleep, and wake throughout the recording sessions.

**Figure 1.**
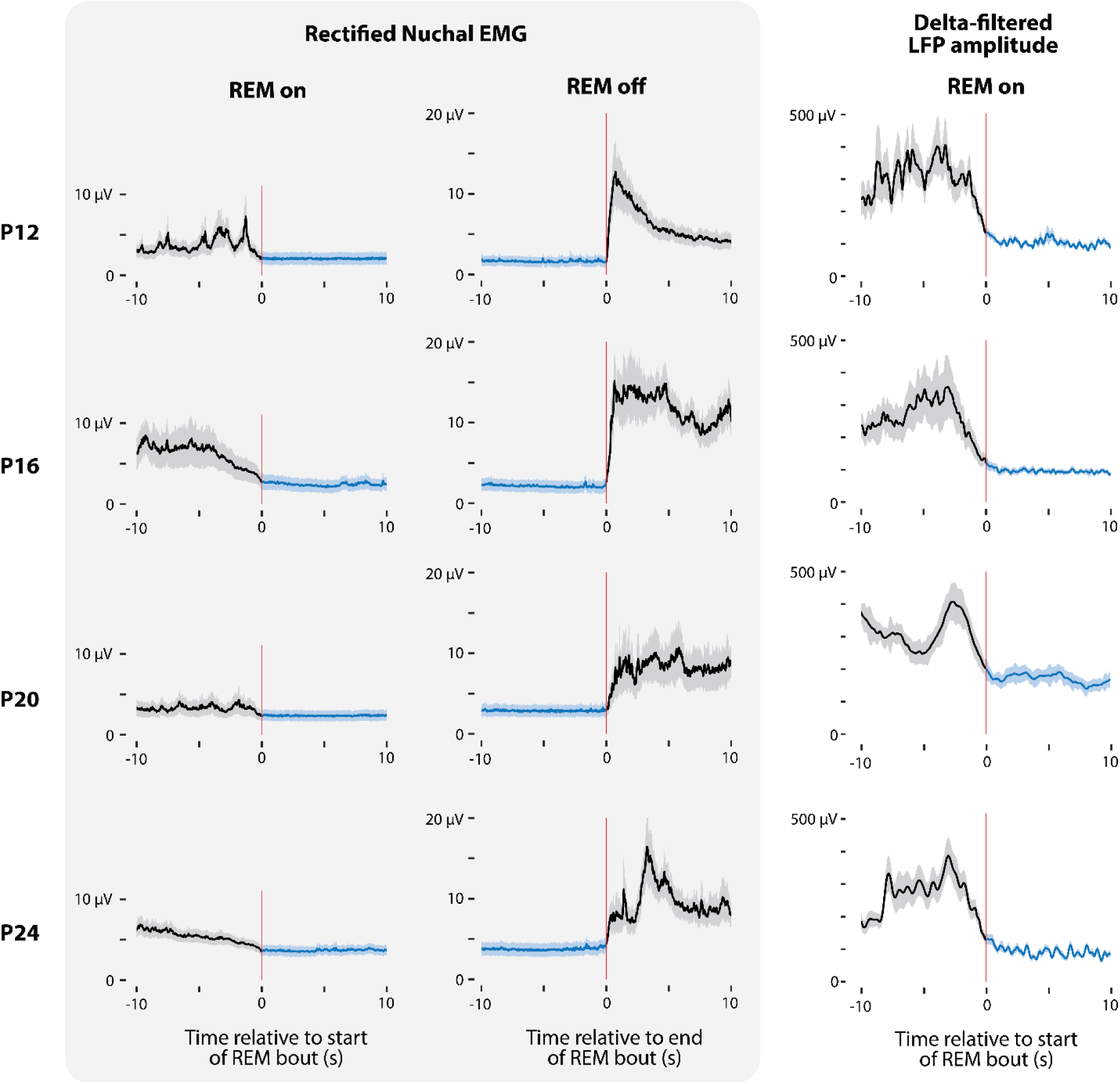
Changes in nuchal EMG and delta power at the onset and offset of REM sleep across development. Left: Rectified nuchal EMG aligned to the start and end of REM bouts at P12, P16, P20, and P24. Each trace shows mean ± SE across all bouts. Blue shading indicates the REM period, with time 0 marking the transition into/out of REM sleep. Right: Same as left, but for delta-filtered (0.5–4 Hz) LFP power aligned to REM onset, averaged across animals.

**Figure 2** *left* shows the increased delta power (0.5-4 Hz) of NREM sleep relative to quiet wake. At P16, P20, and P24, NREM sleep in M1 also showed an increase in alpha (8-15 Hz) power, likely reflecting the developmental emergence of sleep spindles, which in humans appear in the first 2 postnatal months (Robert, 1982 p.198; Sokoloff et al., 2021). REM sleep showed a markedly different oscillatory profile than NREM sleep.

**Figure 2.**
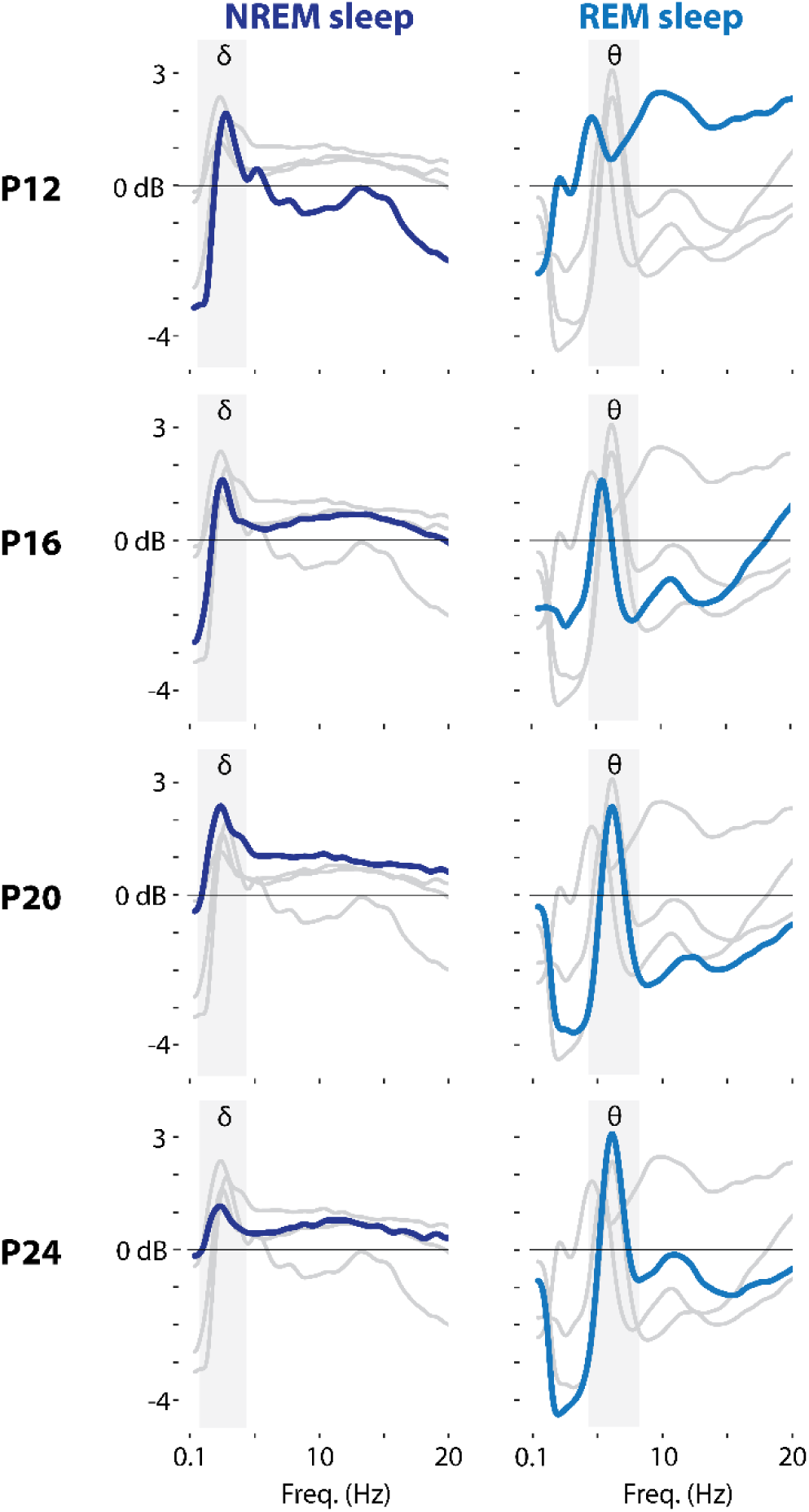
Normalized LFP power spectra during NREM and REM sleep across development. Decibel-converted LFP power spectra (0.1 to 20 Hz) from M1 during NREM (dark blue) and REM (blue) sleep at P12, P16, P20, and P24 (top to bottom). For each age, spectra were normalized to power during quiet wake and then averaged across animals. The colored line represents the mean power for that age and behavioral state, whereas the gray background lines show the corresponding spectra for the other three ages. Frequency bands corresponding to delta (δ) and theta (θ) are indicated. For non–decibel-normalized (raw) power spectra, see **Figure S1**.

Relative to periods of quiet wake, REM sleep in P12 pups was associated with a general increase in power across all frequencies. At P16, P20, and P24 pups, periods of REM sleep showed an increase in power restricted to the theta band (4-8 Hz), along with a decrease in power in the delta and alpha bands (**Figure 2** **right**). REM sleep was also associated with an increase in slow gamma power. The developmental emergence of prominent theta power in M1 between P12 and P16 was also evident in the raw power spectrum; increased power in the theta band was not present at P12, but was present at P16, P20, and P24 (**Figure S1**).

## Increasing delay from REM onset to first twitch

Representative recordings of the onset of REM sleep at each age are shown in **Figure 3**, illustrating the defining electrophysiological and behavioral features of the state across development. Each panel shows the characteristic drop in nuchal EMG tone and reduction in delta power in the LFP at REM onset. From P16 onward, an approximately 6 Hz theta band oscillation is also visible. Myoclonic twitches were readily observed during REM sleep at all ages, although their temporal distribution within each bout changed markedly with age.

**Figure 3.**
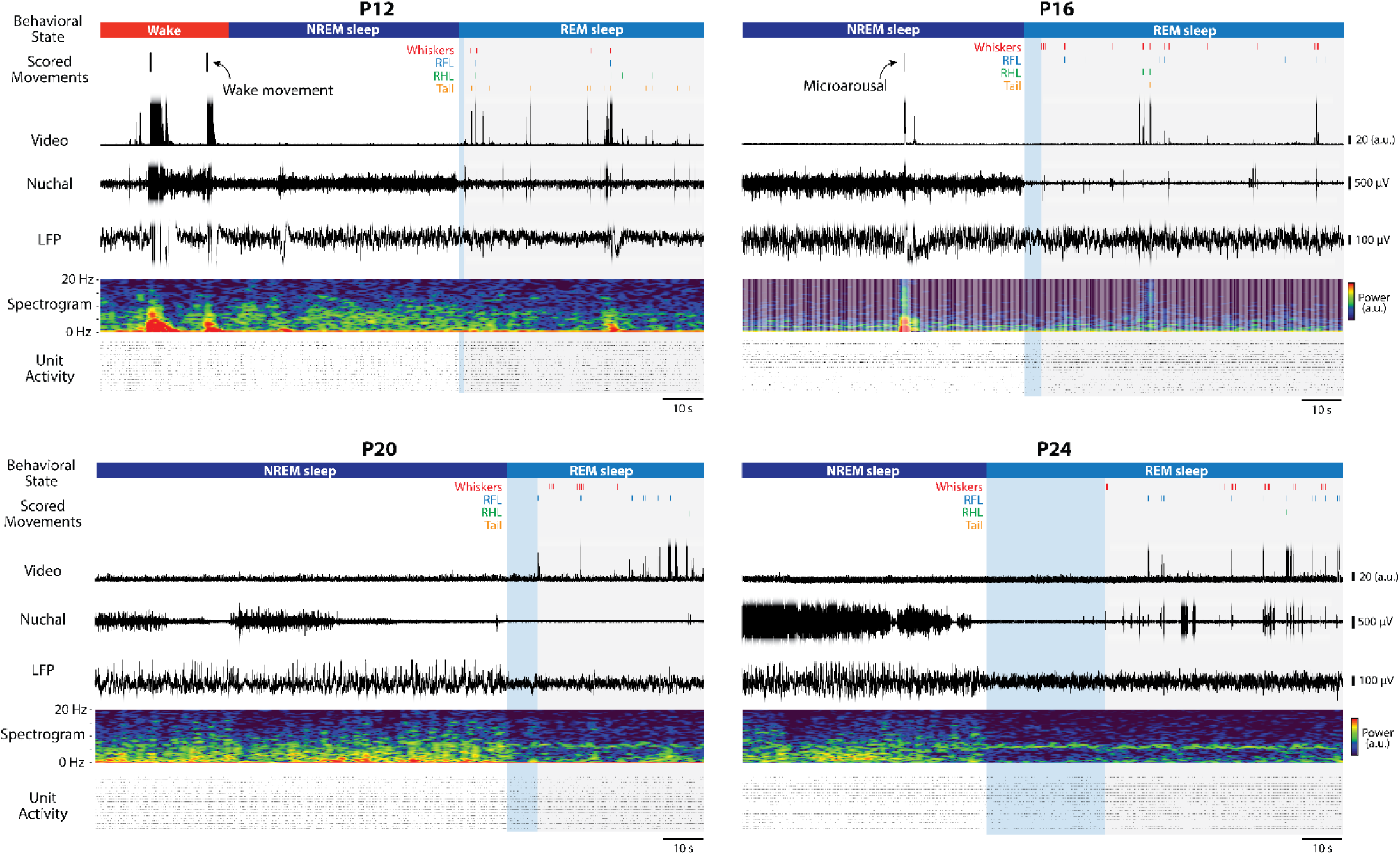
Representative recordings fromM1 across development showing transitions from NREM to REM sleep. Example 200-s segments from P12, P16, P20, and P24 rats illustrating transitions from NREM to REM sleep. From top to bottom within each age, the following signals are shown: scored behavioral state, scored movements (wake movements, arousals, and twitches), ROI-based video analysis of movement, nuchal EMG, M1 LFP trace, M1 LFP spectrogram (0–20 Hz), and spiking activity of isolated M1 units. Light blue shading indicates the interval from REM onset to the first behaviorally scored twitch.

In P12 pups, twitches occurred almost immediately following the onset of REM sleep, often continuing throughout the bout. By contrast, at P20 and P24, the first twitch was delayed, producing an extended period of atonia without twitches—resembling tonic REM in older animals. These representative traces suggest a developmental shift in the structure of REM sleep, from bouts dominated by phasic activity at early ages to longer bouts containing a clear tonic phase preceding clusters of twitches.

To quantify the change in twitch timing across development, every REM bout was plotted as a raster of twitches, ordered by bout duration (**Figure 4A**). Each tick mark represents a scored twitch from any body part. At P12, twitches were distributed relatively uniformly throughout each REM bout, producing a dense, continuous pattern. By P16, however, the onset of each REM bout was often marked by a pause in twitching, followed by a return to regular twitch activity. This trend became more pronounced at P20 and P24, where the early portion of each bout was dominated by extended periods of atonia with little or no twitching, followed by a progressive increase in twitch rate later in the bout. The mean twitch rate for each age group (red dashed line in **Figure 4A**) highlights this developmental progression, with the highest early-bout twitch density at P12 and a clear ramping pattern by P20 and P24.

**Figure 4.**
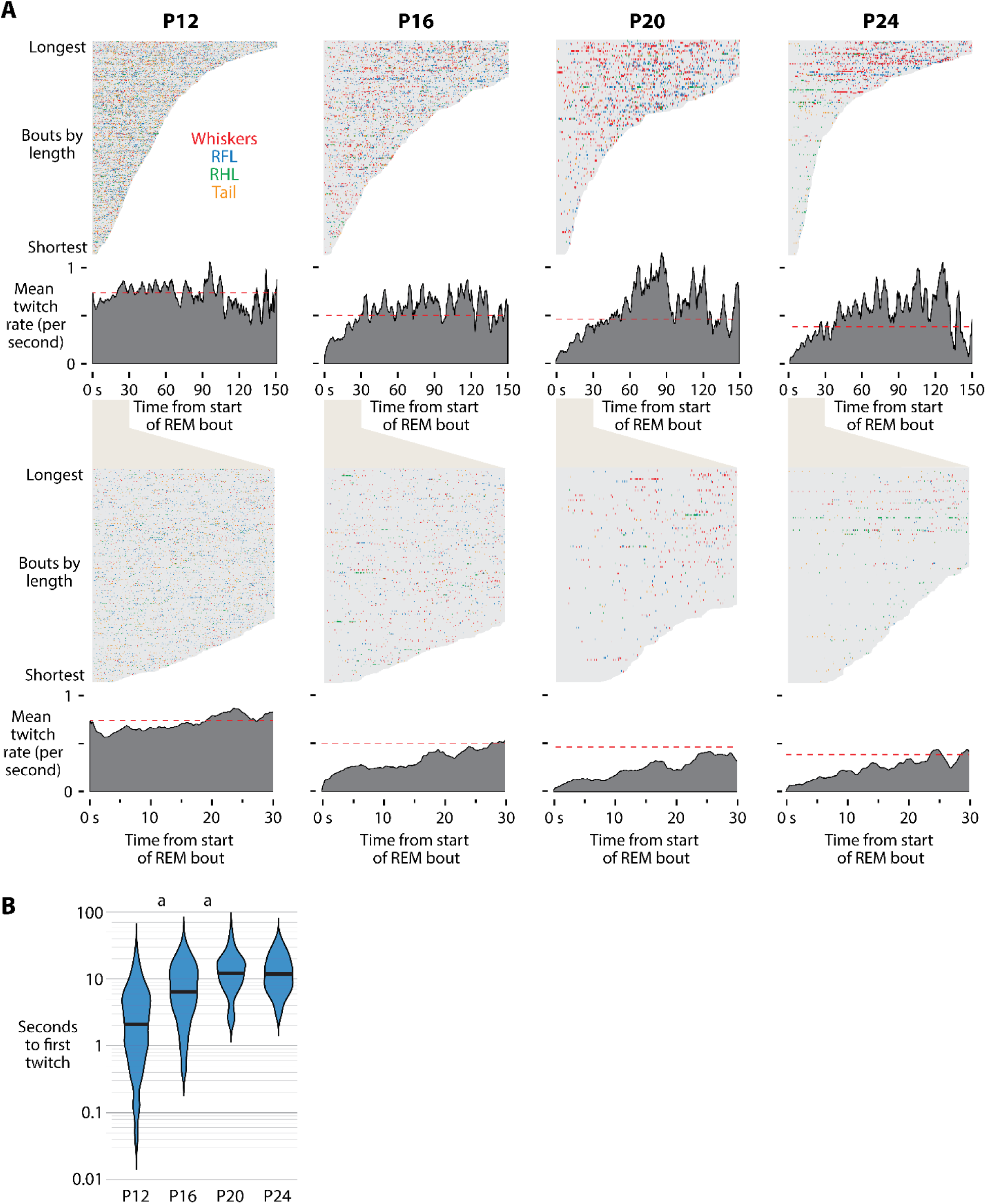
Temporal structure of twitching within REM sleep bouts across development. A *Top:* Each panel shows up to the first 150 s of all REM sleep bouts at P12, P16, P20, and P24 (left to right), sorted by duration from longest to shortest. The shaded gray region represents the duration of each bout, and individual twitches are plotted as colored ticks over each bout. Below, the dark gray trace shows the mean twitch rate across all bouts for each second, with the red dotted line indicating the mean twitch rate across the entire REM bout at that age. *Bottom:* Same data as above but zoomed to the first 30 s of each bout, illustrating the reduced twitch rate at REM onset at P16–P24. B Violin plots show the time from REM onset to the first twitch for all bouts at each age. The letter “a” above P12 and P16 denote a significant difference (p < 0.0125) from the following age.

To quantify this change, we measured the latency from REM onset to the first twitch in each bout (**Figure 4B**). Median latency to the first twitch increased systematically with age from P12 to P20 (P12: 2.1 s; P16: 6.5 s; P20: 12.3 s; P24: 12.0 s). A one-way Kruskal-Wallis test revealed a significant main effect of age on latency to first twitch (H(3) = 343.3, P<0.0001), with post-hoc comparisons showing significant increases from P12 to P16 and again from P16 to P20. These results confirm a developmental shift in REM sleep architecture from bouts dominated by phasic activity to those containing a distinct tonic phase preceding the onset of twitching.

## Developmental differentiation of tonic and phasic REM sleep

To determine whether this developmental difference in the latency to the first twitch captured broader developmental changes in REM sleep structure, REM bouts were subdivided based on the timing of twitches. Periods within ±2 s of a twitch were classified as phasic REM, and all remaining periods of REM sleep were tonic REM (**Figure 5A**).

**Figure 5.**
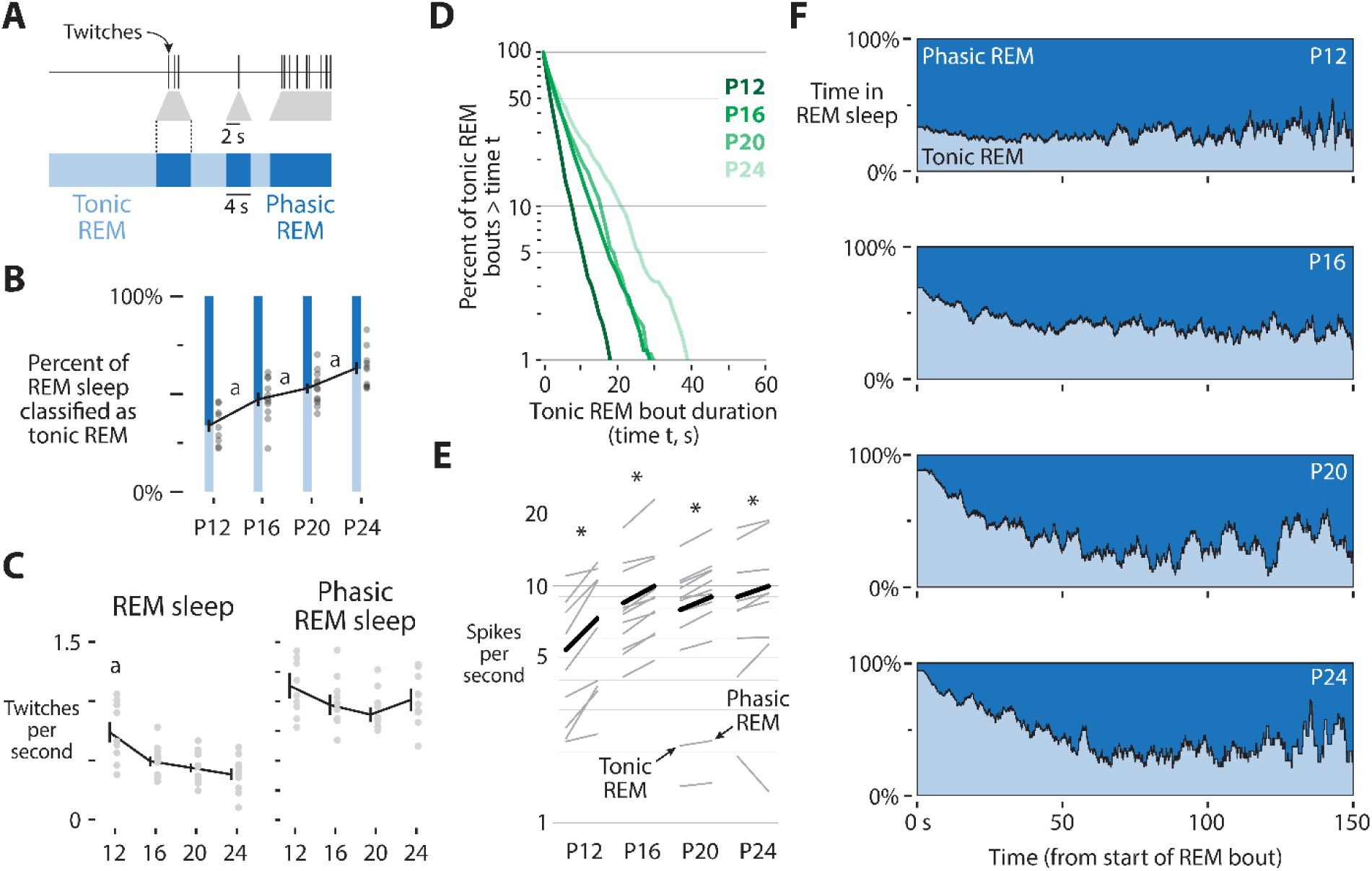
Developmental emergence of tonic and phasic REM sleep. **A** Illustration of the criteria used to distinguish tonic and phasic REM sleep. Phasic REM was defined as any REM period occurring within ±2 s of a behaviorally scored twitch; all remaining REM periods were classified as tonic REM. **B** Percentage of total REM sleep classified as tonic REM across ages. The letter “a” indicates a significant difference (*p* < 0.0125) from the following developmental age. C Twitch rate (twitches/s) during all REM (left) and phasic REM (right). Lines show mean ± SE; gray dots represent individual animals. Letters denote significant age differences as in B. D Percent of tonic REM bouts longer than time t by age, showing that periods of tonic REM increase in duration from P12 to P24. E Mean M1 spiking rate (spikes/s) during tonic and phasic REM at each age. Gray bars denote individual animal means; black bars denote the overall mean of that age. At all ages, M1 neurons fire at a higher rate during phasic REM than during tonic REM (*p* < 0.015). F Likelihood of being in tonic (light blue) or phasic (blue) REM for each second of a REM bout (0–150 s). At P12, the likelihood is relatively stable, whereas by P16—and especially by P20 and P24—tonic REM dominates early in the bout before progressively decreasing.

At each age, pups spent a smaller percentage of time in phasic REM and a larger percentage of time in tonic REM (**Figure 5B**). Tonic REM occupied 34.1% of total REM at P12, 47.6% at P16, 53.2% at P20, and 63.6% at P24. A one-way ANOVA revealed a significant effect of age on the percentage of tonic REM (F(3,38) = 14.8, *p* < 0.0001). Post-hoc comparisons show a steady increase in tonic REM across age, with each age differing significantly from the preceding age.

We also observed that the overall twitch rate during REM sleep decreased significantly with age (**Figure 5C**; F(3,38) = 8.00, *p* < 0.0005), reflecting the overall reduction in phasic activity. However, within phasic REM periods, the twitch rate remained stable across ages (F(3,38) = 1.32, *p* = 0.28), indicating that while the overall twitch rate decreased across development, its rate within phasic episodes did not change.

We also observed a significant age-related difference in the duration of tonic REM bouts, with bouts being shortest at P12 and longest at P24 (**Figure 5D**; logrank test, χ^2^ = 475.72, df = 3, p < 0.0001). At P12, only 5.7% of tonic REM bouts were 10 seconds or longer, whereas at P24 this proportion increased to 27.9%. Further, although tonic REM bouts were distributed relatively uniformly across the REM bout at P12, by P16 they were much more likely to occur at the start of the bout (**Figure 5F**). This tendency for REM bouts to begin in tonic REM strengthened further at P20 and P24, with 80% of bouts starting in tonic REM, mirroring the typical sequence seen in adult rodents (Bueno-Junior et al., 2023; Boscher et al., 2024)

Neural activity in M1 also differed systematically between tonic and phasic REM (**Figure 5E**). At every age, single-unit firing rates were significantly higher during phasic than tonic REM (P12: t(8) = -3.9, *p* < 0.005; P16: t(10) = -3.62, *p* < 0.005; P20: t(11) = -4.76, *p* < 0.0001; P24: t(9) = -3.21, *p* < 0.01), consistent with prior reports that phasic REM is accompanied by increased cortical activity (McCarley and Hobson, 1970; Bueno-Junior et al., 2023). Together, these findings suggest that by the end of the third postnatal week, REM sleep in rats is differentiated into alternating tonic and phasic substates that resemble those described in adult mammals (Brankačk et al., 2012; Bueno-Junior et al., 2023).

## Oscillatory differences between tonic and phasic REM

Spectral analyses of M1 LFPs revealed that tonic and phasic REM differ not only in motor activity but also in their oscillatory composition (**Figure 6**). At P12, phasic REM was characterized by a broadband increase in power relative to quiet wake, whereas tonic REM showed a uniform decrease in power below ∼20 Hz. This broad increase in power during phasic REM is likely the result of event-related potentials produced by the long duration response to twitches at this age (Dooley and Blumberg, 2018; Reid et al., 2025). Beginning at P16, both tonic and phasic REM exhibited an increase in power at gamma (30-55 Hz) and fast gamma (65-100 Hz) frequencies compared to quiet wake (Brankačk et al., 2012). Consistent with prior reports, phasic REM showed significantly higher power than tonic REM in both gamma bands (Simor et al., 2016; Avigdor et al., 2025).

**Figure 6.**
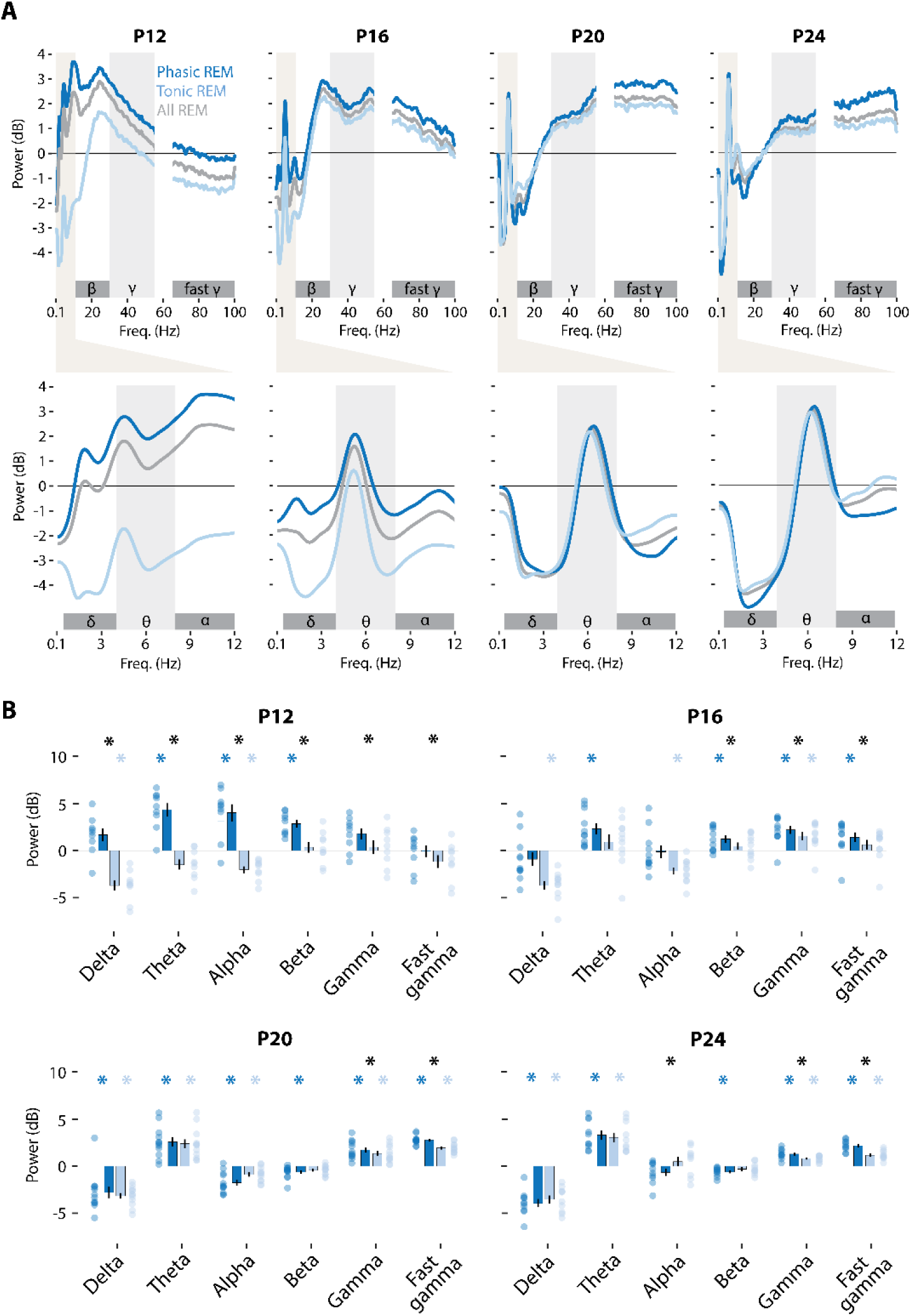
Spectral properties of tonic and phasic REM sleep across development. **A** Decibel-converted LFP power spectra from M1 during REM sleep (gray), tonic REM (light blue), and phasic REM (dark blue) from P12 to P24 (left to right). As in **figure 2**, power was normalized to quiet wake. Power is shown from 0.1–100 Hz, with a magnified view of 0.1–15 Hz below. Frequency bands used for delta (0.5–4 Hz), theta (4–8 Hz), alpha (8–12 Hz), beta (12–30 Hz), gamma (30–55 Hz), and fast gamma (65–100 Hz) are indicated. The 55–65 Hz range (notch filter) was excluded from analysis. Raw power data can be found in **Figure S2.** B Mean power (±SE) for each frequency band during tonic and phasic REM at each age. Individual animal values are shown as light blue (tonic) and dark blue (phasic) dots. A dark blue asterisk indicates that phasic REM power significantly differs from 0 (p < 0.0083), a light blue asterisk indicates that tonic REM power significantly differs from 0 (p < 0.0083), and a black asterisk indicates a significant difference between tonic and phasic REM (p < 0.0083).

Beginning at P16, a prominent theta peak (4–8 Hz) emerged in both tonic and phasic REM, increasing in strength at each subsequent age. The prominence of this theta band is particularly evident at P20 and P24, where relative to quiet wake the theta peak is accompanied by a dip in the frequencies on either side (**Figure 6A**). To assess whether theta frequency differed between REM substates within each age, we performed paired *t*-tests at P16, P20, and P24 (**Figure S2B**). At P16, theta frequency did not differ significantly between phasic and tonic REM (*t*(10) = 0.48, *p* = 0.64). By P20, theta frequency was significantly higher during phasic REM (6.08 ± 0.08 Hz) than tonic REM (5.80 ± 0.08 Hz; *t*(11) = 8.2, *p* < 0.0001). A similar pattern was present at P24, where phasic REM exhibited faster theta (6.09 ± 0.06 Hz) than tonic REM (5.91 ± 0.07 Hz; *t*(9) = 3.00, *p* = 0.015). Together, these comparisons indicate that reliable substate differences in theta frequency emerge by P20 and are maintained at P24 that is consistent with prior work in adult rodents (Vanderwolf and Robinson, 1981; Bueno-Junior et al., 2023).

Finally, at P24, we found a significant difference between tonic and phasic REM in the alpha band (t(9) = -3.50, p < 0.0001), illustrating an increase in power in the alpha band during tonic REM. Notably, across ages and frequency bands, whenever a significant difference between substates was detected, phasic REM almost universally exhibited greater power than tonic REM—with the sole exception of this alpha band effect at P24. An increase in alpha power during tonic REM was also observed at P20, though not significant (t(11) = -3.11, p = 0.091). This increase in alpha power during tonic REM is consistent with prior research (Simor et al., 2016, 2019). Together, these results highlight a developmental trajectory toward the substate-specific oscillatory patterns seen in adults, marking the emergence of distinct tonic and phasic REM dynamics by P24.

## Discussion

The tonic and phasic REM substates described here did not appear fully formed, but instead emerged gradually, similar to previous reports describing the gradual emergence of REM and NREM sleep across infancy (Seelke and Blumberg, 2008; Blumberg et al., 2020). At P12, the rate of twitches was the same throughout a REM bout. But by P16, the twitch rate was much lower at the start of a REM bout, and the latency to the first twitch increased (**Figures 3 and 4**). This difference became even more pronounced at P20 and P24. Inspired by this, we separated REM sleep into tonic and phasic periods. Phasic REM consistently exhibited elevated cortical activity and gamma power in the LFP (**Figure 5**), while tonic REM developed a fast-alpha component in the LFP by P24 (**Figure 6**). Importantly, twitch rate within phasic periods remained constant across ages, even as overall twitching declined (**Figure 5C**). This suggests that the REM sleep of early infancy aligns most closely with phasic REM, with tonic REM emerging later as a distinct substate whose defining physiological features first develop between P16 and P24. Together, these findings define the developmental onset of REM substates and link their behavioral and neural oscillatory features across early postnatal development.

## Increasing complexity of sleep states across development

Sleep in early life is dynamic, reflecting, among other things, the ongoing development of neural circuits. Through the first postnatal week of life, cortical neurons exhibit intermittent, sensory-driven bursts of activity separated by periods of silence (Khazipov et al., 2004). While there are state-dependent differences in the frequency of these bursts (Dooley et al., 2020), the bursts of activity occur across both sleep and wake. Thus, neural activity alone is not sufficient to define behavioral state, necessitating reliance on peripheral measures like muscle tone and movement. At this age, rat pups cycle almost exclusively between wakefulness and REM sleep, which is defined by atonia punctuated by myoclonic twitches of the limbs, whiskers, and tail (**Figure 7**). Because of the prominence of twitching, behaviorally, REM sleep in infancy resembles phasic REM in adults. However, there is one important distinction: early REM is marked by increased sensory responsiveness—particularly when compared with wake, when sensory feedback is actively suppressed (Mukherjee et al., 2017; Dooley and Blumberg, 2018; Murata and Colonnese, 2018)—whereas adult phasic REM is characterized by high arousal thresholds and reduced sensory responsiveness (Ermis et al., 2010; Brankačk et al., 2012).

**Figure 7.**
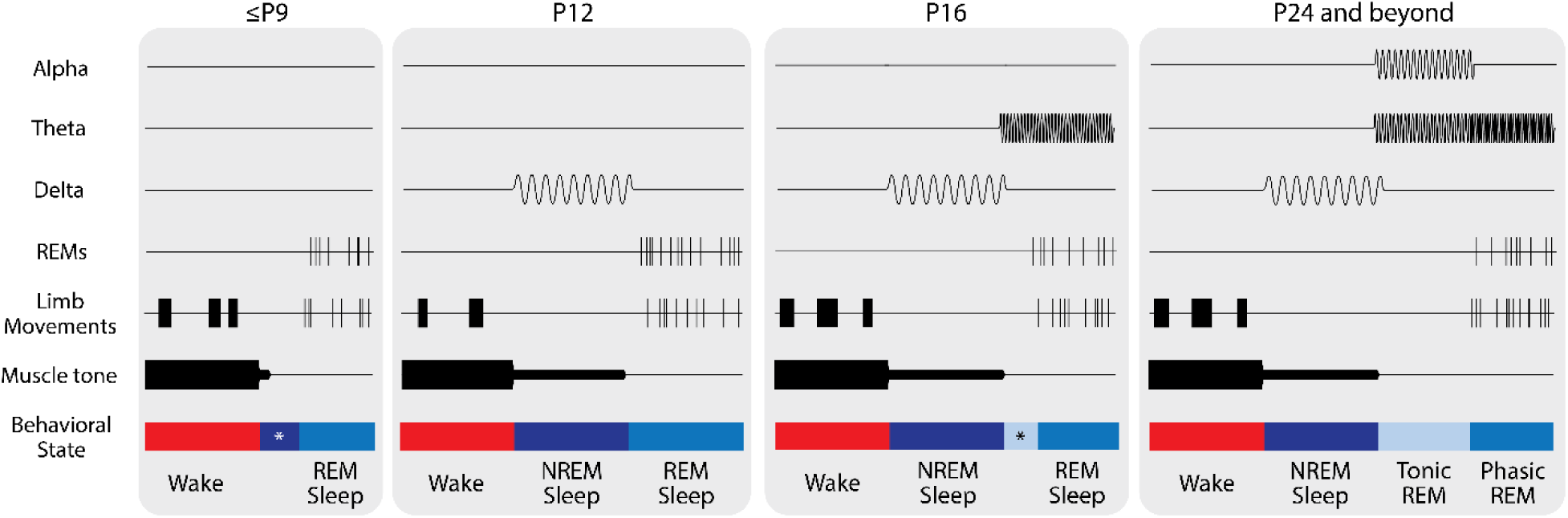
Developmental progression of sleep state organization. Schematic summary illustrating the development of the behavioral and neural features of sleep from ≤P9 through P24 and beyond. From left to right, panels depict characteristic features of each state at representative ages, including muscle tone, limb movements, and cortical oscillations (delta, theta, and alpha bands). At ≤P9, sleep consists primarily of wake and active (REM) sleep, with early signs of NREM-like activity emerging at the beginning of sleep bouts. By P12, cortical delta clearly delineates NREM sleep. At P16, the delayed onset of the first twitch following REM onset marks the behavioral emergence of tonic REM, though without any distinct cortical oscillations between tonic and phasic REM. By P24, tonic and phasic REM are fully differentiated, distinguished by their theta frequency, alpha power, twitch rate, and neural activity profiles. Asterisks denote ages at which some—but not all—features of a given state are present.

By one week of age, a second, more quiescent sleep state, begins to appear. In week-old pups, a period where twitches are suppressed, respiration is regular, and muscle tone remains low but not absent begins to emerge at the transition from wake to sleep (Blumberg et al., 2005; Seelke and Blumberg, 2008). This state—often called quiet sleep—does not yet have all the features of NREM sleep, as cortical activity remains discontinuous and characteristic delta oscillations are not yet present. From P10–P12, the corticothalamic networks necessary to support synchronized delta oscillations develop, providing the first positive electrophysiological signature of NREM sleep (Seelke and Blumberg, 2008). From this point forward, pups reliably alternate between NREM and REM sleep, marking the onset of a stable two-state sleep cycle.

The present investigation builds upon this developmental sequence by identifying another point of differentiation—this time within REM sleep itself. Although tonic REM can be defined behaviorally as the twitch-free periods of REM sleep at any age, by P16 we observed a consistent bout structure, with twitch-free tonic REM reliably occurring at the onset of a REM bout. This organization strengthened with age: by P24, tonic REM was not only more likely to initiate a REM bout (**Figure 5F**) but also persisted for significantly longer bouts (**Figure 5D**). Between P16 and P24, these behavioral distinctions corresponded closely to emerging oscillatory differences that parallel those seen in adults: phasic REM maintained high levels of neural activity, elevated gamma power, and faster theta oscillations, whereas tonic REM gradually developed a distinct fast-alpha oscillation by P24.

Thus, as cortical circuits continue to develop, sleep becomes organized at multiple hierarchical levels—first into REM and NREM states, and then into tonic and phasic substates within REM. Over this period, the proportion of time spent in tonic REM increases, from 34% at P12 to 64% at P24 (**Figure 5B**), results broadly consistent with the decrease in phasic REM previously described across these ages (Vogel et al., 2000). Given estimates that over 90% of REM sleep is tonic REM in adult rodents (Brankačk et al., 2012), this trend likely continues beyond P24. Yet despite the growing prevalence of tonic REM, the rate of twitching within phasic episodes remains stable from P12 to P24.

This persistence suggests that, even as phasic REM occupies a smaller share of total sleep, its defining behavioral and neural features are preserved—consistent with a conserved role in sensorimotor integration and plasticity across development.

Together, these findings position the emergence of tonic and phasic REM as the latest refinement in a developmental continuum of sleep organization—beginning with the active sleep of the neonate and culminating in the complex, layered architecture of the juvenile and adult brain.

## Functional roles of phasic and tonic REM

REM sleep, and its substates, are accompanied by muscle atonia. As summarized in **Table 1**, tonic and phasic REM represent distinct but complementary modes of coordinated activity in the brain and the body. Phasic REM is defined by the appearance of twitches and rapid eye movements, accompanied by bursts of cortical activity and elevated gamma and theta power. This coincides with high arousal thresholds and reduced sensory responsiveness, indicative of a brain deeply engaged in internally-generated patterns of activity. In contrast, tonic REM is characterized by the absence of movements and increased alpha power—features that more closely resemble relaxed wakefulness. Specifically, theta oscillations are slower, cortical activity is lower, and sensory responsiveness is partially restored, suggesting a partial re-engagement with the external world. In adults, REM bouts typically begin in the tonic state and transition into phasic periods, reflecting a shift from externally receptive to internally driven modes of processing.

The presence of a state that, behaviorally, resembles phasic REM from the earliest postnatal days—and its persistence across development—strongly supports the idea that this state serves a fundamental developmental function. Myoclonic twitches, the defining feature of phasic REM, are expressed against a backdrop of muscle atonia and generate discrete bursts of sensory feedback that reverberate throughout the somatosensory and motor systems. These self-generated movements provide a rich stream of temporally precise neural activity that helps to construct and refine sensorimotor representations (Blumberg et al., 2022). Thus, phasic REM produces twitches that act as a training signal for the developing nervous system, driving correlated neural activity across distributed sensorimotor circuits. In doing so, this activity helps to shape sensory maps (Dooley and Blumberg, 2018; Gómez et al., 2021, 2023) and calibrate internal models of movement (Dooley et al., 2021). As development proceeds, however, REM sleep begins to include longer periods of behavioral quiescence—suggesting the emergence of a distinct, yet complementary, substate.

If phasic REM provides the scaffold for building sensorimotor representations, what might be the function of the activity driven by the later-emerging tonic REM? One clue lies in its defining behavioral feature—the near-complete absence of movement. Unlike phasic REM, tonic REM is marked by behavioral quiescence and, in humans, increased sensory responses driven by the partial re-engagement of thalamocortical sensory pathways (Wehrle et al., 2007). These observations suggest that the two substates promote distinct modes of neural coordination: phasic REM strengthens activity patterns that arise from self-generated movements, whereas tonic REM organizes patterns of activity that occur in the absence of movement—more akin to the sensory attention or contextual processing typically seen during wake.

Consistent with this distinction, dream studies report that individuals awakened during phasic REM describe vivid, movement-rich scenes, whereas those awakened during tonic REM describe more static or observational imagery (Berger, 1968; Wehrle et al., 2007). Of course, such reports are inherently subjective, but they nonetheless provide a framework for linking behavioral quiescence, sensory responsiveness, and network activity. Viewed developmentally, the delayed appearance of tonic REM suggests that linking self-generated movements to sensory feedback may be a prerequisite for later representations of sensory input independent of movement.

In this framework, the emergence of tonic REM around the end of the third postnatal week may represent more than a change in oscillatory patterning—it signals the onset of coordinated cortical activity capable of sustaining internally-generated sensory representations in the absence of movement. Phasic REM continues to provide the building blocks of embodied experience through sensorimotor reactivation, while tonic REM introduces a state in which these newly established circuits can be stabilized, synchronized, and integrated across cortical areas. Together, these complementary modes of REM sleep may provide the foundation for the increasingly complex forms of behavior emerging during the transition from infancy to juvenile life. By parsing REM sleep into its component substates, we can begin to trace how sleep’s functions expand in parallel with development.

## Limitations and future directions

Several limitations of the present study warrant consideration. First, recordings were performed in head-fixed animals. Although pups readily entered sleep and displayed clear behavioral and electrophysiological signatures of REM sleep, immobilization likely influenced bout structure, particularly at older ages where REM episodes were often shorter than those described in freely moving rodents (e.g. Boyce et al., 2016; Bueno-Junior et al., 2023).

Second, we did not assess arousal thresholds, which in adult animals differ markedly between phasic and tonic REM and could offer an additional functional metric of substate differentiation during development. However, to date no rodent studies have directly examined differences in arousal thresholds across tonic and phasic REM (see **Table 1**). Notably, one recent study (Boscher et al., 2024) performed patch clamp recordings on thalamic somatosensory neurons during REM sleep, inferring stronger inhibition of somatosensory inputs during tonic REM, and weaker somatosensory inhibition during phasic REM. Although these results appear contradictory, all human studies have assessed auditory thresholds whereas the rodent findings reflect somatosensory inhibition. The discrepancy may therefore arise from differences between sensory modalities rather than conflicting conclusions about REM substates.

Nevertheless, these constraints highlight important next steps. The present results provide a foundation for future studies using freely moving preparations and physiological measures of arousal using multiple modalities to more fully characterize how REM sleep and its substates change across early development and into adulthood. Such approaches will be critical for linking behavioral state transitions to their underlying neural dynamics and for testing how substate organization contributes to the emergence of sleep–wake architecture.

## Conclusions

Taken together, our findings show that REM sleep begins as a largely undifferentiated state. At P12, the youngest age examined here, REM bouts are punctuated by brief twitch-free periods, but these lack the defining features of tonic REM and do not exhibit the predictable sequence—from tonic to phasic REM—seen later in development. Between P16 and P24, we observed the emergence of a reliable, behaviorally distinct substate: tonic REM bouts appear at the onset of most REM bouts, lengthen with age, and acquire increasingly specific oscillatory signatures, including a fast-alpha rhythm by P24. In parallel, phasic REM maintains its characteristic high neural activity, gamma elevation, and—by P20—faster theta oscillations. By the end of the third postnatal week, REM sleep is therefore composed of two clearly differentiated substates whose behavioral and electrophysiological profiles closely resemble tonic and phasic REM in adults. This progression marks a key developmental refinement in sleep architecture, transforming the early, twitch-dominated REM of infancy into the structured microarchitecture characteristic of the adult brain. The developmental onset of tonic REM may therefore mark a transition toward representational capacities that rely less on moment-to-moment sensorimotor input and more on internally coordinated cortical dynamics.

## Acknowledgments

We thank Mark Blumberg, Greta Sokoloff, Nick Sattler and Madi Reid for feedback on earlier versions of this manuscript. This research was supported by the Sleep Research Society Foundation’s Career Development Award to J.C.D.

## Author contributions

J.C.D. designed research; J.C.D. performed research; J.C.D. data curation; J.P.K and J.C.D. analyzed data; J.P.K and J.C.D. visualization; J.C.D writing – original draft; J.P.K and J.C.D. writing – review & editing; J.C.D. supervision; J.C.D. resources

