## Supplemental Figures for "The developmental emergence of tonic and phasic REM sleep in rats"

**Figure S1**

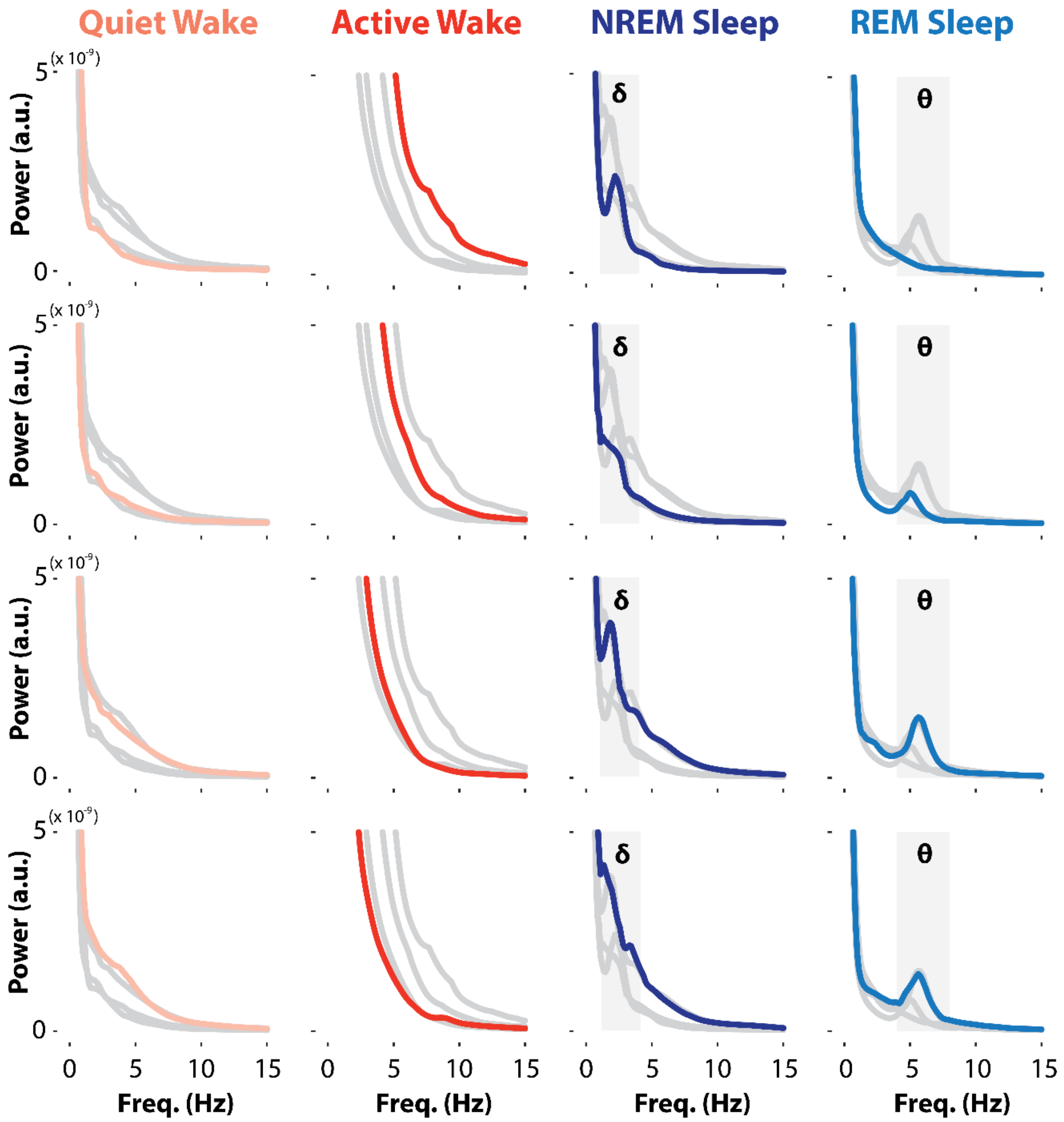


**Figure S1. Developmental changes in the raw power spectrum across behavioral states**

Raw power spectra (0–15 Hz) for each age (P12 at the top, P24 at the bottom) and each of the four behavioral states (quiet wake, active wake, NREM sleep, REM sleep, from left to right). In each panel, the focal age and behavioral state are shown in color (consistent with the color scheme used in the main text), whereas data from other ages in the same behavioral state are shown in gray. Note the prominent peak in the delta range (1–4 Hz) during NREM sleep and the growing peak in the theta range (4–8 Hz) from P16 onwards during REM sleep.

**Figure S2**

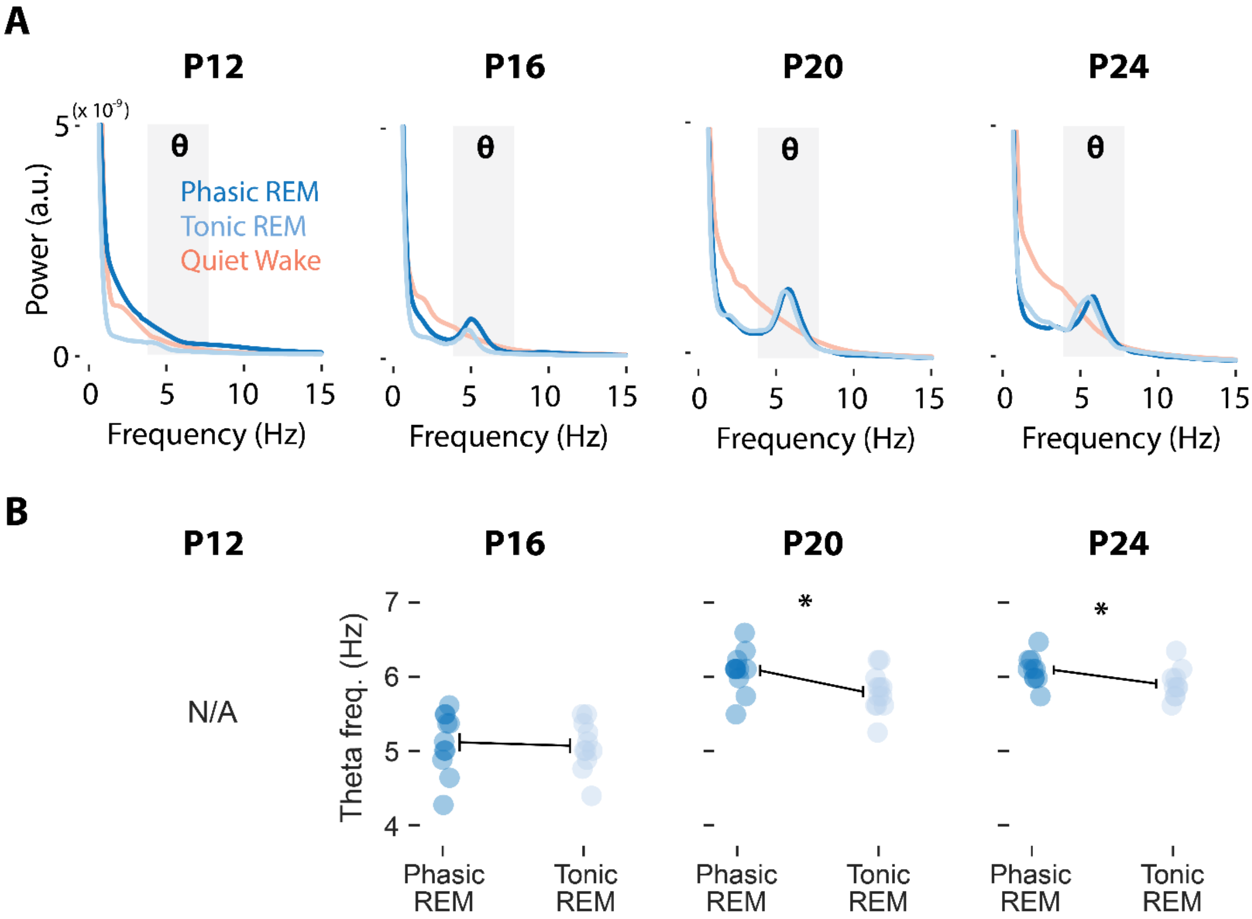


**Figure S2. Spectral features of tonic and phasic REM sleep across development.** **A** Raw power spectra (0–15 Hz) for phasic REM (dark blue), tonic REM (light blue), and quiet wake (light red) at each age (P12, P16, P20, and P24; left to right). **B** Peak theta frequency (Hz) for phasic and tonic REM at each age (P16, P20, P24). Because a theta peak was not present at P12, that age is omitted (see **A** and **Figure 6A**). Within each age, paired *t*-tests assessed differences between phasic and tonic REM. Theta frequency did not differ between substates at P16, but was significantly higher during phasic REM at both P20 and P24.
